# Removal of epicuticular wax has no effect on the rate of cuticular wax deposition, which is constant in mature cherry laurel leaves

**DOI:** 10.1101/2024.11.27.625630

**Authors:** Jitka Janová, Tereza Kalistová, Jiří Kubásek

**Affiliations:** Department of Experimental Plant Biology, Faculty of Science, University of South Bohemia, Branišovská 1760/31, České Budějovice, Czech Republic

**Keywords:** Plant cuticle, wax regeneration, ^13^C labelling, gas exchange, photosynthesis

## Abstract

- The plant cuticle, multifunctional hydrophobic air-to-plant boundary, vary dramatically among plant species and responds dynamically to environmental and biotic factors. These dynamics, certainly driven by cost-benefit selection, are poorly understood up to now.
- ^13^C labelling and compound-specific isotope analysis open a new avenue to study the cuticle dynamics. We studied side specific leaf cuticular wax regeneration, in different compounds, and separately for intra-(IW) and epicuticular wax (EW), as well as the effect of epicuticular wax removal on the regeneration rate.
- Mature leaves that reached the final area before the start of the experiment deposited new IW at first, but quickly equilibrated with EW. Removal of EW accelerated this equilibration but not the net rate of new wax deposition. The n-alkanes had the fastest turnover. Pentacyclic triterpenoids (ursolic acid) had a surprisingly slow turnover due to the large pool but also slow deposition rate. Adaxial and abaxial leaf surfaces exhibited small but consistent differences.
- We demonstrated for the first time that EW removal does not affect wax deposition rate in evergreen leaf cuticle. We also confirmed that collodion discriminates between EW and IW well and does not inhibit future leaf metabolism and wax deposition in our species.

## Introduction

The plant cuticle is a thin extracellular layer produced by the plant epidermis covering all aboveground living plant tissues (Riederer & Muller, 2006) and also the root tips (Berhin *et al*., 2019). Its main function is to protect against water loss, (Schönherr & Schmidt, 1979; Burghardt & Riederer, 2003), but it also reflects/absorbs UV radiation (Krauss *et al*., 1997), protects from pathogen attack (Aragón *et al*., 2017; Kalistová & Janda, 2023), and helps keeping the plant surfaces dry and clean due to its hydrophobic nature, so called “lotus effect”, respectively (Barthlott & Neinhuis, 1997).

Structurally, the cuticle can be divided into two components: Cuticle matrix (MX) and cuticular waxes. The MX consists of an insoluble polymer usually formed by cutin and a minor fraction of polysacharides (as a consequence of the cell wall-cuticle continuum), amino acids, phenols and other compounds tightly bound to the polymer (Philippe *et al*., 2020). Waxes, on the contrary, are soluble and consist typically of a mixture of linear molecules with a long carbon chain (18C to 54C) and various functional groups, mainly acids, alkanes, aldehydes, ketones, primary and secondary alcohols and esters (Samuels *et al*., 2008; Yeats & Rose, 2013). Moreover, pentacyclic triterpenoids (PCT) and small amounts of aromatic compounds may also be present (Samuels *et al*., 2008; Jetter *et al*., 2018). The amount and composition of the wax vary across plant species, the organs, ecotypes, and growing conditions, with coverage in the range from less than one to more than 1000 μg · cm^-2^ (Jetter *et al*., 2018). Depending on the localisation, waxes are recognised as intracuticular wax (IW), embedded in the cutin polymer, and epicuticular (or extracuticular) wax (EW) deposited on the surface (Buschhaus & Jetter, 2011). The waxes form the main barrier against water loss. The wax fraction can be extracted by immersion of leaves or isolated cuticles into an organic solvent such as chloroform or hexane. Wax removal increases the permeability of the cuticular membrane to water vapour by an average of two to three orders of magnitude (Schreiber & Schonherr, 2009). It was shown that exclusively the IW fraction is responsible for gas barrier function in most species (Zeisler & Schreiber, 2016; Zeisler-Diehl *et al*., 2018). There are some evidence that cuticular presence of PCTs render both, IW and EW important for this function (Jetter & Riederer, 2016). Zeisler-Diehl et. al, however, used also plant species with cuticular triterpenoids and confirmed their previous conclusion that only IW renders substantial resistance to water loss (Zeisler-Diehl *et al*., 2018).

In this study, the dynamics of both, aliphatic compounds and PCTs, are shown. PCTs are considered to be of medical and healthcare importance and plant cuticles are studied as a potent source of various biologically active triterpenoids (Szakiel *et al*., 2012). Numerous triterpenoid derivatives are present in many leaf and fruit cuticles (Buschhaus & Jetter, 2011) and may exceed other compounds in their coverage. In cherry laurel plant, PCT ursolic acid was found in high concentration in the IW (Jetter & Schäffer, 2001a). The function of triterpenoids was studied mainly on the fruit cuticle. They do not seem to contribute to the water barrier properties (Seufert *et al*., 2022), but they probably provide a protection from harmful pests and pathogenic microbes, modify the mechanical toughness of the fruit peel and maintain their postharvest quality (Fang & Xiao, 2021). In addition, PCT seem to enhance the mechanical strength of the fruit cuticle by functioning as nano-fillers (Tsubaki *et al*., 2013). Functional studies, however, are lacking in leaf cuticles.

The cuticular matrix (MX) is synthesized early in leaf development, the cuticle precursor, the procuticle, covers the very earliest epidermal cells in shoot apices and leaf primordia. It is an electron-dense layer approximately 20 nm thick visible with transmission electron microscopy (Jeffree, 2018). MX of expanded leaves is not renewed for the rest of leaf longevity(Riederer & Schönherr, 1988; Kubásek *et al*., 2023). In contrast, cuticular wax composition varies during leaf ontogeny, as was shown for *Prunus laurocerasus* (Jetter & Schäffer, 2001b), *Hedera helix* (Hauke & Schreiber, 1998) or *Fagus sylvatica* (Prasad & Gülz, 1990). This developmental changes are not only due to selective wax loss/interconversion, but also due to the synthesis and deposition of new compounds, even in mature evergreen leaves (Kubásek *et al*., 2023). However, it is not yet known whether the removal of EW affects the synthesis/deposition of new wax.

Isotope labelling technique is a useful method to study the dynamics and potential regeneration of cuticular wax. First approach is to irrigate the plants by heavy water (D_2_O) and to detect deuterium in the new wax (Kahmen *et al*., 2011; Gao *et al*., 2012). In combination with compound specific isotopic analysis, it is possible to follow “the fate” of individual wax compounds. Gao et al. (2012) determined the wax renewal half-time in timothy, *Phleum pratense*, to be 2-3 days for C16 and C18 fatty acids (probably not of the cuticular origin, see later), 5-16 days for waxes with carbon chain lengths C22-C26, and 71-128 days for the very long chain (VLC) component of waxes (C27 to C31). The composition of the newly synthesized waxes may change in response to the ontogenetic leaf development(Jetter & Schäffer, 2001a) or seasonal and environmental conditions (Kerfourn & Garrec, 1992).

The second method, pulse-chase labelling by ^13^CO_2_, was applied to plant cuticle by (Kubásek *et al*., 2023). We let the plants assimilate air with highly ^13^C enriched CO_2_ for a short period (≈ one hour) and traced the carbon newly incorporated into the cuticle compounds with a time resolution of hours to several weeks.

Cherry laurel (*Prunus laurocerasus, Rosaceae*) is one of the most popular models for cuticle studies (together with *Arabidopsis thaliana*). Its leaf cuticle is relatively thick (2 to 8 µm), easily enzymatically isolable, and the leaves are hypostomatous with stomata exclusively on the abaxial leaf side, allowing a direct comparison of astomatous and stomatous cuticles. Many studies on this species characterised cuticle and cuticular waxes (Stammitti *et al*., 1996), changes in wax chemical composition during the ontogeny (Jetter *et al*., 2000; Jetter & Schäffer, 2001a), cuticle’s permeability (Schreiber *et al*., 2001), location of pentacyclic triterpenoids exclusively in intracuticular wax layer (Jetter *et al*., 2000) or nanoscale heterogeneity in the cuticle (Perkins *et al*., 2005; Diarte *et al*., 2021) in last three decades. On the other hand, only limited information about wax dynamics (Jetter & Schäffer, 2001a) and virtually nothing about de-novo wax synthesis and cuticle regeneration after disturbance are available up to now. Not only for cherry laurel but also in general.

In this study we tested whether: i) the cuticular wax is synthetized and deposited (in IW and EW) in mature, fully developed leaves of *Prunus laurocerasus*, ii) the removal of EW affects the rate of new wax deposition and iii) regeneration dynamics differ between EW and IW and in different compounds. We also made comparison with annual plant species measured in our pilot experiments.

## Material and Methods

### Plant material

This study was performed on four commercially obtained cherry laurel plants (*Prunus laurocerasus L*., *Rosaceae*, var. Novita) ca. 40 cm tall. Plants in pots with mass-produced soil were kept in a greenhouse with natural photoperiod for two months before the start of the experiment. Maximum day temperature of 26°C and minimum night temperature of 16°C were maintained, and relative humidity ranged from 50-80 %.

Ten mature leaves were selected at each plant and equipped with labels for subsequent sampling (see ^13^CO_2_ labelling and sampling strategy). The outlines of each leaf were carefully drawn on the paper to see if the leaves expanded between the labelling and sampling (see Sfig. 1). Epicuticular wax (EW) was removed from one half of each leaf by applying collodion (Fluka), a solution of 4-8% nitrocellulose in diethyl ether:ethanol (1:1, v/v), in a thin layer with a soft brush (Zeisler & Schreiber, 2016). After evaporation of the solvent (≈ one min), the collodion films along with the adhering epicuticular wax were stripped off with a tweezer and collected in a glass vial. The adaxial and abaxial leaf sides were treated sequentially and the strips kept separately and labelled as epicuticular wax-free (**EWF**). The other half of the leaf remained intact, native, and labelled as control (**Cont**.). Variants were separated by a central vein and this experiment design allowed us to compare treatments and controls from a single leaf, i.e. without the influence of leaf age or position (**Fig. 1**).

**Fig. 1.**
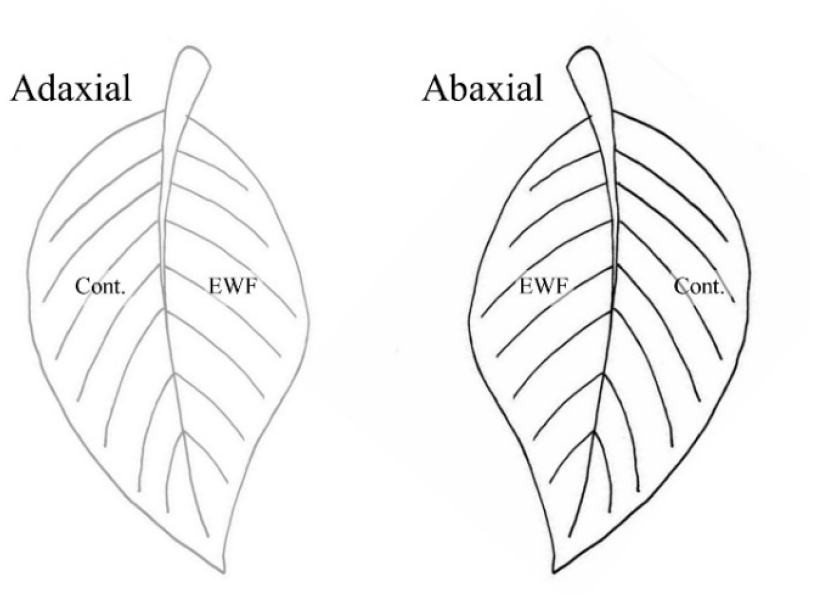
Design of wax sampling. Each leaf selected for wax sampling was divided into two halves, separately for adaxial and abaxial leaf side. One leaf half was collodion-sampled before labelling and again in particular time after labelling. Because initial collodion treatment removed substantial load of epicuticular wax (EW) this treatment is denoted as ‘epicuticular wax free’ (EWF). Contrary to, control half of leaf was sampled only once in particular time after labelling, thus unreduced EW coverage was subjected to ^13^C enrichment during chase period (1h to four weeks).

### Cryo-Scanning Electron Microscopy (Cryo-SEM)

A fresh leaf samples serving as control or collodion treated fresh leaves with removed epicuticular waxes were glued to the aluminium target using Tissue-Tek® O.C.T. Compound (EMS) and frozen in liquid nitrogen. Then they were transferred into the high vacuum cryo-preparation chamber (CryoALTO 2500, Gatan, UK). In the cryo-preparation chamber under the vacuum was the sample sublimated for 3 minutes at the temperature −90 °C and coated with 5 nm gold. The sample was observed using a Field Emission Scanning Electron Microscope (JSM-7401-F, JEOL) pre-cooled at -135 °C. Images were obtained at an accelerating voltage in the range 1-4 kV.

### ^13^CO_2_ Labelling and sampling strategy

The pulse labelling with heavy stable carbon isotope (^13^C) was done according Kubásek et al. (2023) in August 2023. One cherry laurel plant was placed in a gas-tight Plexiglas box (internal dimension 60 × 60 × 60 cm and volume 190 l) for one hour. The box was illuminated by linear LED sources with total PPFD of ca. 400 μmol m^-2^ s^-1^, the temperature was kept at 20°C. A fan was placed inside the box to homogenize the air and reduce the resistance of leaf boundary layer. After equilibrating the air in the room and in the box, the box was hermetically sealed and 80 ml of ^13^CO_2_ (> 99 % atoms, Sigma-Aldrich) was injected through a septum in the box at the beginning and again in the middle (after 30 min) of the labelling, yielding ca. 800 μmol CO_2_ mol^-1^. Previous experiment with natural CO_2_ confirmed (data not shown), that CO_2_ concentration was not exhausted to less than 400 μmol CO_2_ mol^-1^ at the end of the labelling. After labelling the plants were returned to the greenhouse for the rest of the experiment.

Leaves were harvested before the start of the labeling (t = 0 h; only several samples to obtain natural ^13^C abundances), immediately thereafter (t = 1 h) and then after 12, 24, 48, 96, 168 (1 wk.), 336 (2 wks.), 504 (3 wks.) and 672 h (4 wks.). Leaves were scanned, and the area was analyzed using ImageJ software.

At each time-point, the adaxial and abaxial side of the control (Cont.) and pre-treated (EWF) leaf section (four treatments in total) were harvested separately in four biological replicates. After isolation of EW by collodion (as described in *Plant material* section), 3 to 4 discs of 2 cm diameter, depending on the leaf size, (area of 3.14 cm^2^) were cut to isolate the cuticle.

### Isolation of epicuticular and intracuticular waxes

EWs were striped out from the leaf blade surface along with the collodion film (as described above) and dissolved in two ml of chloroform (VWR Chemicals) on a roller overnight. Chloroform is a widely used solvent for extracting cuticular wax (Riederer & Schneider, 1989) without dissolving the nitrocellulose film. To each sample, 20 μg of *n*-tetracosane (C24 alkane) was added as an internal standard. The samples were transferred into 2 ml analytical glass vials, evaporated to one ml to be prepared for the GC-IRMS analysis.

EW-free leaf discs were immersed in a 2% enzymatic solution of cellulase and pectinase (Schönherr & Riederer, 1986) to isolate the cuticle. The isolation process was accelerated by infiltrating the solution into the discs under vacuum and keeping the samples at 35°C.

Isolated cuticles were divided into astomatous adaxial and stomatous abaxial ones (using a microscope). IW from each cuticle were isolated by immersion in one ml of chloroform with 20 μg of *n*-tetracosane (internal standard) in 2-ml vials at room temperature on the roller overnight. The wax-free cuticular matrices were removed from the vials subsequently.

### Gas chromatography, mass spectrometry and stable isotope analyses

All chloroform extracts were derivatized (50 μl BSTFA; Macherey-Nagel; and 100 μl pyridine; Sigma Aldrich; 2 h at 80°C) to transform OH group of alcohols and acids into the corresponding trimethylsilyl-ethers and –esters (Hauke & Schreiber, 1998). Subsequently the samples were analysed on a gas chromatograph (Trace 1310; Thermo, Bremen, Germany) equipped with a ZEBRON ZB-1 column (30 m × 0.25 mm × 0.25 μm film thickness). One μL (EW) or two μL (IW) of a sample was injected at 300°C in a splitless mode at 1.5 mL min^-1^ column flow (for 1.5 min) continued by split flow at 100 mL min^-1^ for another 1 min and 5 mL min^-1^ for the rest of the analysis. The oven temperature was set at 50°C during injection and next 2 min, followed by a temperature ramp with increase 40 °C min^-1^ up to 185 °C, 20 °C min^-1^ to 275 °C, 4°C min^- 1^ to 300 °C, 10 °C min^-1^ to 320 °C and isothermal for the rest of the analysis (18 min). Individual separated compounds were oxidized to CO_2_ via IsoLink II interphase (Thermo, Bremen, Germany) at 1000°C and introduced into a continuous flow isotope ratio MS (Delta V Advantage; Thermo, Bremen, Germany) to obtain δ^13^C value. Kovats retention indices (RI) were calculated using n-alkane C8 to C40 mixture (Sigma Aldrich) and n-alkanes identified. N- aldehydes were identified comparing RI with values obtained in our previous study where compounds were directly identified using ESI-TOF Mass spectrometry (Kubásek *et al*., 2023).

### Calculations and statistics

^13^C enrichment is expressed as a ^13^C excess. This variable is simply difference between ^13^C abundance in particular sample/compound and its natural abundance (in At% of ^13^C). Natural ^13^C abundance was found to be 1.065 ± 0.001 At% for C29 alkane, 1.063 ± 0.001 At% for C31 alkane and 1.070 ± 0.002 At% for ursolic acid. Thus, natural variability in ^13^C abundance is negligible in our context and led to more than two orders of magnitude lower uncertainty of initial ^13^C content (‘baseline’, about 0.001 At% of dominant alkanes) than the range induced due to the labelling (up to 0.3 At%).

^13^C excess is a nice proxy of newly assimilated carbon turnover. On the other hand, this value results from the mixing model – highly ^13^C enriched carbon added into much larger pool of natural carbon (where ^13^C excess = 0). Total carbon pool of waxes differed substantially (for instance EWF treatment resulted in up to ≈ 80% decrease of subsequently harvested epicuticular wax). Thus, ^13^C excess is not a reliable proxy of newly synthetized/deposited wax in such comparisons.

We calculated the product of particular compound coverage (µg cm^-2^) multiplied by its ^13^C excess (At%) for estimation of newly deposited wax amount, the proxy introduced in Kubásek et al. (2023). Due to previously mentioned negligible variability in natural ^13^C abundance of particular compounds, it seems to be robust measure of newly deposited carbon. The only assumption for the validity of the proxy is that wax compounds are not lost in substantial amounts from the cuticle components (EW and IW, taken separately) during the duration of the experiment (see Discussion).

Finally, we estimated the equilibration rate of newly deposited wax (more precisely its carbon) between IW and EW wax. This calculation relies on the product introduced in previous paragraph, separately for IW and EW.

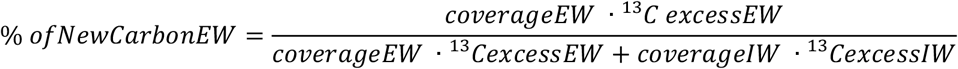

Where coverage is in μg cm^-2^ and ^13^C excess in At%.

Again, this calculation holds true only when the percentage of wax loss due to abrasion/evaporation is negligible or similar between IW and EW and in all treatments. We must be aware particularly for adaxial leaf sides where EW coverage is dramatically lowered after EWF treatment.

Unless other measures are given, we report the mean and standard deviation. T-test (two comparisons) and one-way ANOVA (multiple comparisons within a factor) were used when statistical significance was indicated. Normality and homogeneity of variances were met due to the low variability of the variables of interest compared to their mean. The typical number of replicates is 4 (four independent plants sampled). On the other hand, pooled results are the result of multiple measurements. For example, matrix heatmaps pool adax. and abax. leaf side,s as well as collodion treatments. Thus, N values can be as high as 32 if all values obtained.

## Results

### Leaf area

The leaves were about 30 to 80 cm^2^ in projected area and nicely symmetric. Because the opposite leaf halves were used for two contrasting treatments and main rib was excluded, we calculated the used area as (total area/2)*0.9 (about 10 % of each half accounted for a midrib). These areas did not change during four-week experiment as may be seen in **SFig. 1**.

### Leaf surface before and after collodion treatment

Scanning electron microscopy was employed to observe the effect of collodion before and directly after the collodion-mediated EW removal. It may be seen that not all EW was collected by a one collodion strip, particularly on abaxial leaf side (**Fig. 2h**). It may be the reason for the lower abaxial EW yields in comparison to adaxial EW. On the other hand, a high selectivity of EW removal was reached (yielding the minimum of triterpenoids located only in IW, **STab. 1**). No leaf tissue damage was also observed as the wax continuously regenerated and no necrotic lesions were observed during the experiment.

**Fig. 2.**
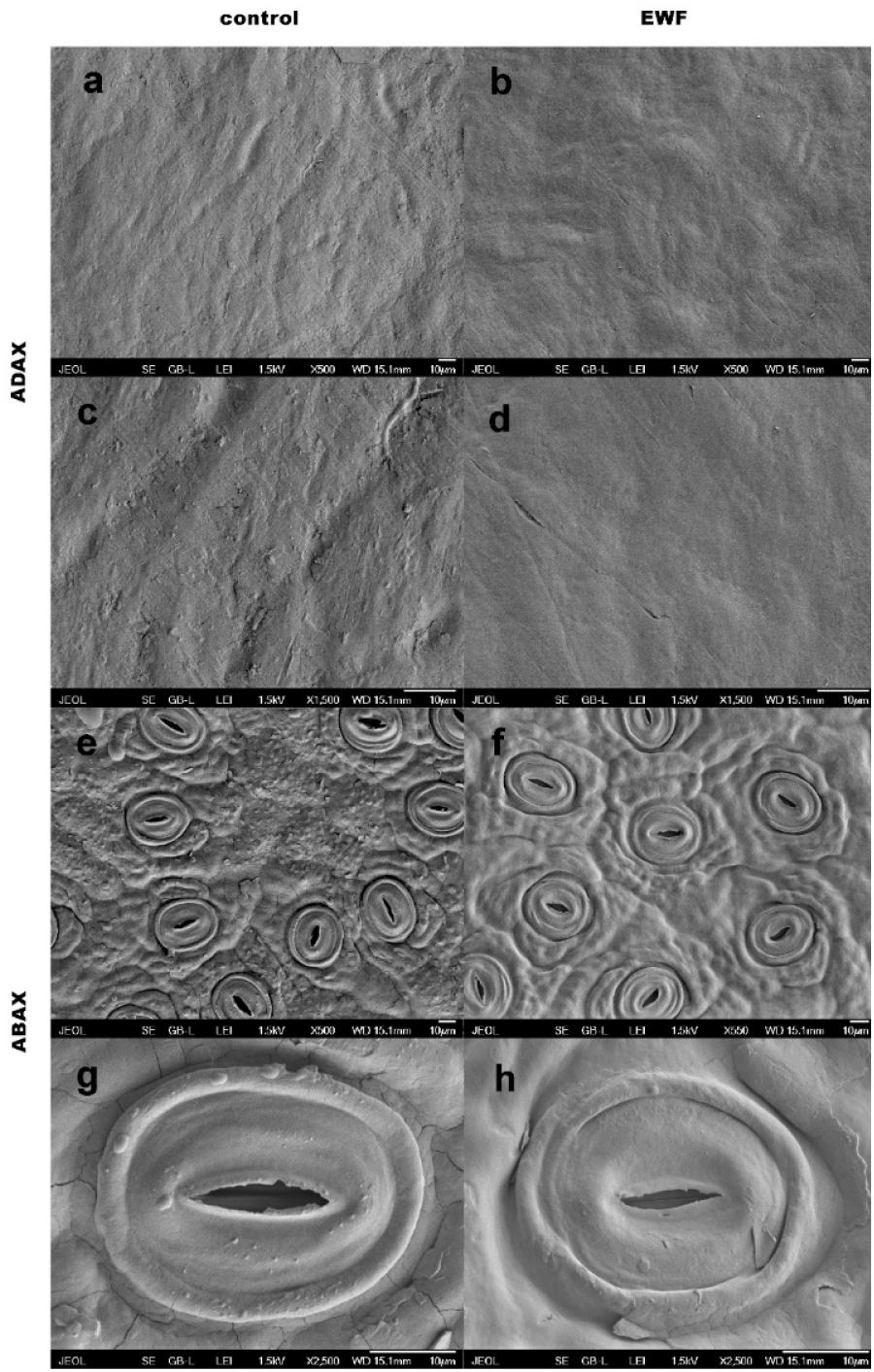
Cryo-SEM images of leaf surfaces of *P. laurocerasus* before (a,c, e, g) and immediately after extraction of EW by collodion (b,d,f,h). The astomatous adaxial (a-d), and stomatous abaxial sides (e-h) are shown in two magnifications.

### Composition of cuticular wax, collodion EW selectivity

Consistent with previous studies, we detected two dominant n-alkanes (nonacosane, C29 and hentriacontane, C31) and two pentacyclic triterpenoids (Oleanolic and Ursolic acids, isomers, both C_30_H_48_O_3_) in *P. laurocerasus* leaf cuticle. Coverage (amount per area, µg cm^-2^) of C29 and C31 alkanes were comparable in EW and IW and consistent during the four-week experiment (**Fig. 3, SFig. 2**). Abaxial leaf side yielded lower EW coverages (C31: 0.48 µg cm^-2^) than adaxial one (C31: 1.13 µg cm^-2^, F(1, 148)=114, p<0.001). The initial EW removal substantially reduced the subsequent EW yield (denoted as ‘epicuticular wax free’, EWF, closed symbols and dotted lines) in contrast to intact leaves (control, open symbols and solid lines), particularly for the adaxial leaf sides and the beginning of the experiment (e.g. ADAX, C31 in time 12 h: 1.1 to 0.22 µg cm^-2^, F(1,6) = 11.4, p=0.015, **Fig. 3b**). However, EWF and control yields, converged towards the end of the experiment (2 to 4 weeks), indicating a new EW deposition (ADAX, C31 in time 648 h: 1.82 to 1.44 µg cm^-2^, F(1,6) = 2.54, p=0.162, **Fig. 3b**). The coverage of IW alkanes was affected by the initial EW removal minimally, but in initial times significantly (ADAX, C31 in time 12h: from 0.85 to 0.65 µg cm^-2^, F(1, 6) =6.24, p=0.047).

**Fig. 3.**
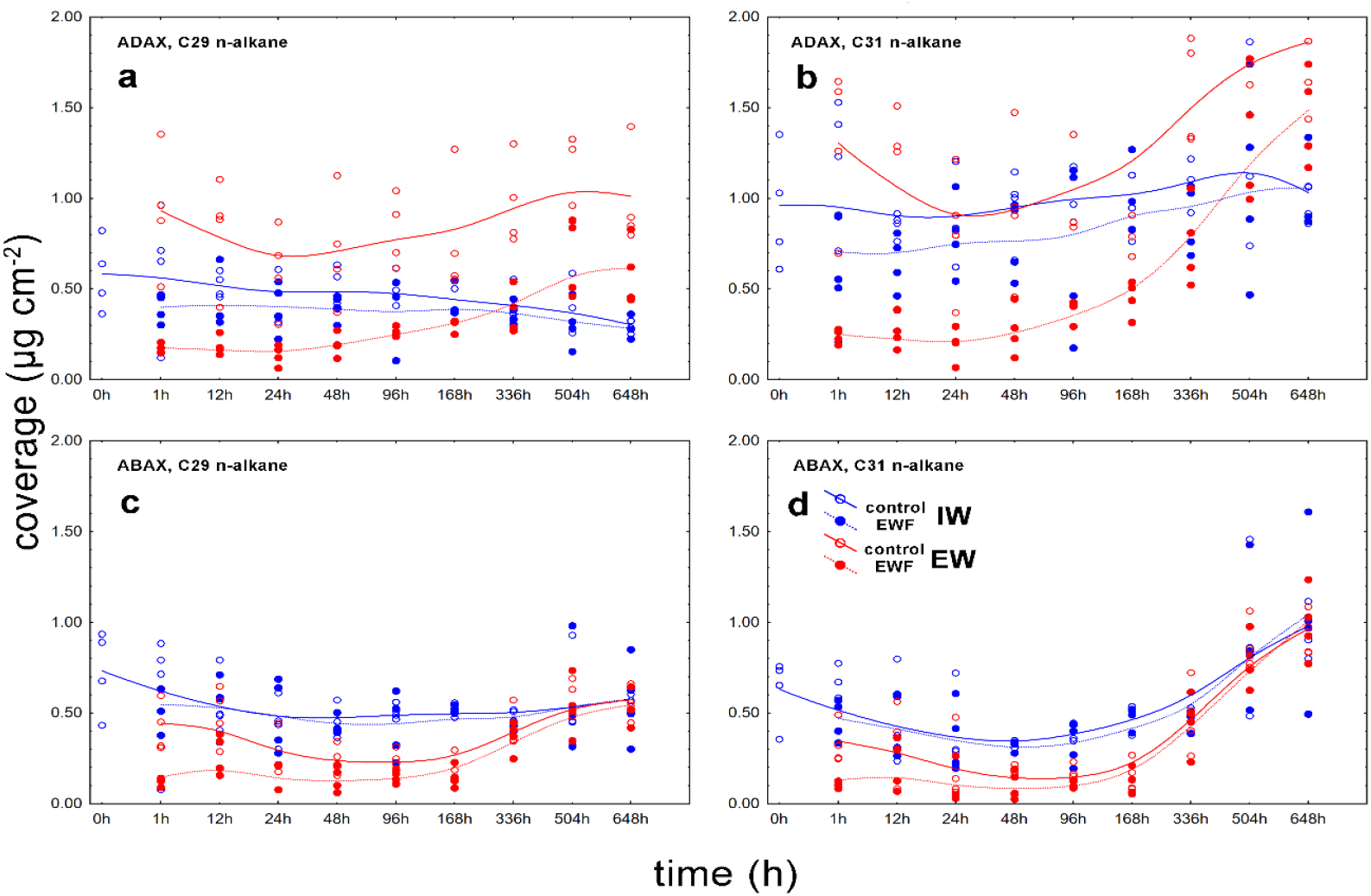
Coverages of the main alkanes (n-nonacosane – a,c; and n-hentriacontane – b,d) present in the *P. laurocerasus* leaf cuticles. EW (red) stands for collodion-collected epicuticular wax, IW (blue) represents for intracuticular wax subsequently extracted from enzymatically isolated cuticles. Open symbols are unaffected leaf halves – controls; closed symbols are leaf halves in which the EW was partially removed by collodion at the beginning of the experiment (‘epicuticular wax free’, EWF). **a, b** show adaxial leaf sides; **c,d** abaxial sides. N=4.

In contrast, the coverages of pentacyclic triterpenoids (mainly ursolic acid) was more than two orders of magnitude higher in IW (mean: 16.74 µg cm^-2^) than in EW (mean: 0.12 µg cm^-2^, F (1, 280)=580, p<0.001, **Fig. 4**), with some differences between treatments and leaf sides (**STab 1**.). Previous research provided strong evidence that pentacyclic triterpenoids are absolutely dominant in IW (and absent or very minor in EW) in *P. laurocerasus* and some other species studied. Thus, any presence of these compounds in EW can largely be considered as an artefact, revealing that the EW collecting method is not selective only for the EW layer.

**Fig. 4.**
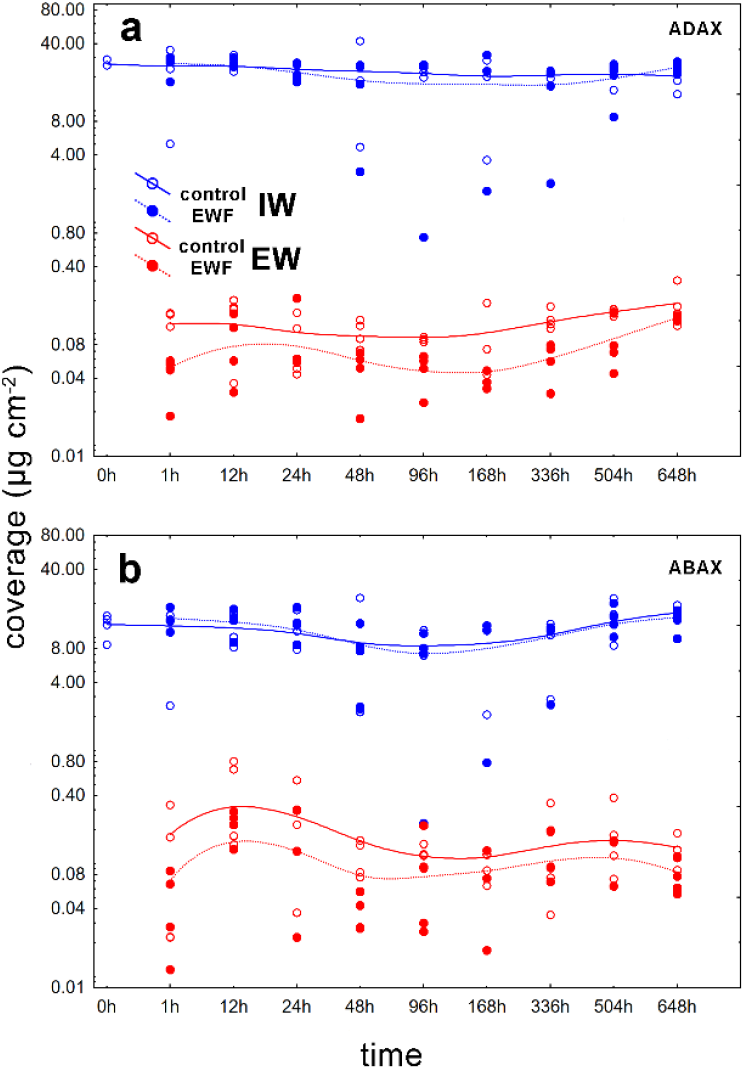
Coverage of ursolic acid in the adaxial (a) and abaxial (b) cuticles of *P. laurocerasus* leaves. EW (red) stands for collodion-collected epicuticular wax, IW (blue) for intracuticular wax subsequently extracted from enzymatically isolated cuticles. Open symbols are unaffected leaf halves - controls, closed symbols are leaf halves in which the EW was partially removed by collodion at the start of the experiment (‘epicuticular wax free’, EWF). Note that y-axis is logarithmic.

Similar to the alkanes, the ursolic acid coverage is not affected in the IW layer and is reduced (although probably artificially) in the EW layer by the EWF treatment. The recovery of EW ursolic acid, in contrast to the alkanes, was not visible during the four-week experiment. Coverages of all detected compounds may be seen in **SFig 2**. (n-alkanes) and **SFig. 3** (other compounds).

### ^13^C enrichment in different compounds and cuticle layers

The most abundantn-alkane, hentriacontane (C31), exhibits a characteristic ^13^C enrichment = excess of ^13^C abundance over natural abundance in course of the experiment (**Fig. 5**), very similar to the second most abundant n-alkane, nonacosane (C29) (**SFig. 4**). IW was enriched faster in all cases. EWF treatment resulted in a faster and higher enrichment in EW (**Fig. 5b,d, SFig. 4b,d)** in contrast to the control treatment (see largely non-overlapping confidence limits). Minor n-alkanes (< C29) were virtually unenriched, except for C33, tritriacontane (**Fig. 6**).

**Fig. 5.**
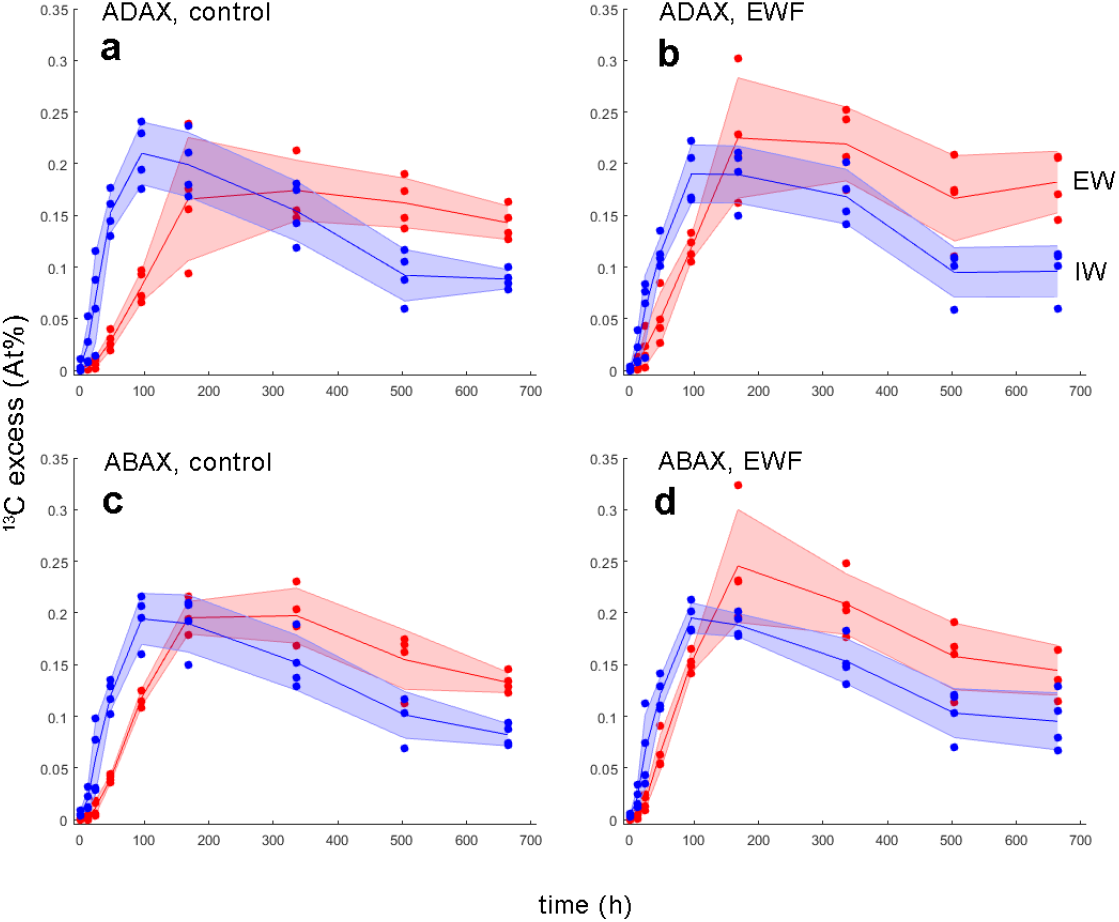
Timecourse of ^13^C excess of n-hentriacontane in the *P. laurocerasus* leaf cuticles. Intracuticular wax (IW, blue) and epicuticular wax (EW, red) are shown separately. **a,b** show adaxial leaf side, **c,d** abaxial leaf side. **a,c** show unaffected leaf halves (control), **b,d** halves in which the epicuticular wax was partially removed by collodion at the start of the experiment (‘epicuticular wax free’, EWF). Lines represent mean, shaded areas are one standard deviation, N=4.

**Fig. 6.**
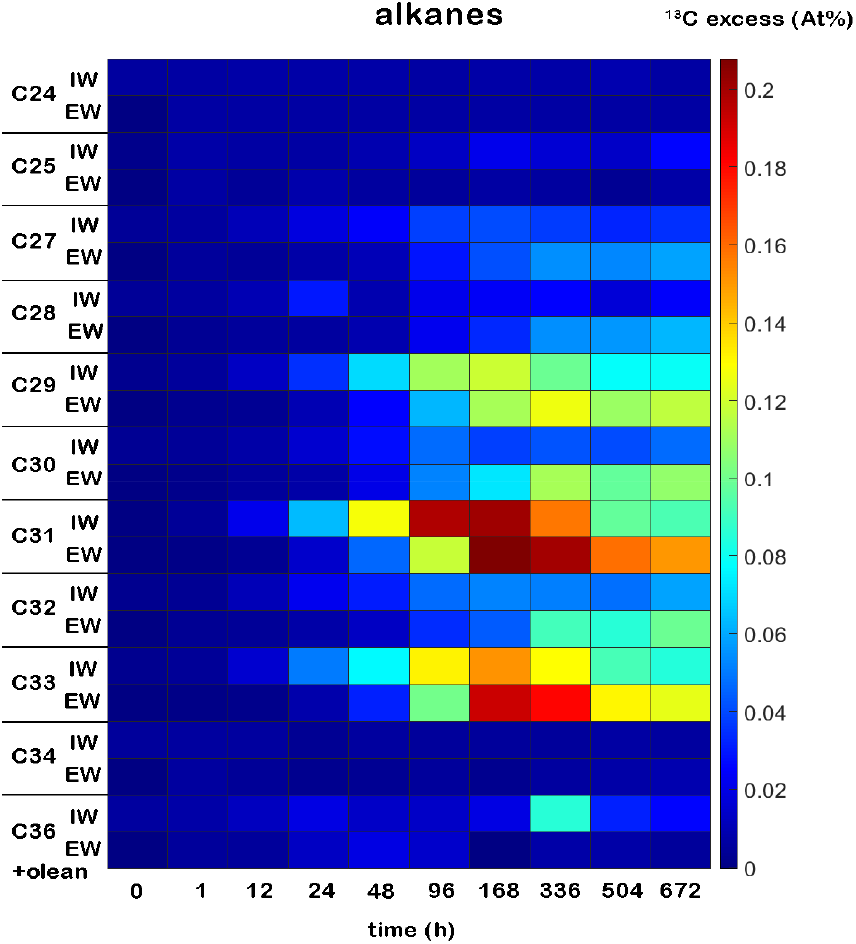
Time course of ^13^C excess of alkanes detected in the *P. laurocerasus* leaf cuticles separately for intra- (IW) and epicuticular wax (EW). Oleanolic acid co-elutes with C36 alkane. Both leaf sides and all treatments are pooled. N=8-32.

Ursolic and oleanolic acids exhibit almost zero ^13^C enrichment in IW, although there seems to be some enrichment in EW (**Fig. 6, Fig. 7, STab. 1**). Oleanolic acid elutes together with C36 alkane and ursolic acid elutes with C34 aldehyde in our set-up. This is a limitation of the compound specific isotope analysis that it separates compounds of interest only by retention time. Since the (1) ursolic acid coverage in EW is low, typically less than one percent of that in IW (see **Fig. 4, STab. 1**), and (2) the co-elution mentioned above exists, an enrichment in co-eluting compounds, may be visible in EW ursolic acid peak. We tried also different GC setup to minimise co-elution in ursolic acid retention time. When four selected EW samples with a particularly high (apparent) ^13^C excess in ursolic acid were analysed on a different column (LION LN-1MS, 30 m × 0.25 mm ID x 0.1 µm film thickness), EW ursolic acid ^13^C excess was negligible and comparable to IW samples, typically < 0.020 At% (data not shown). Thus, even EW samples exhibit almost zero ^13^C enrichment in PCT ursolic acid.

**Fig. 7.**
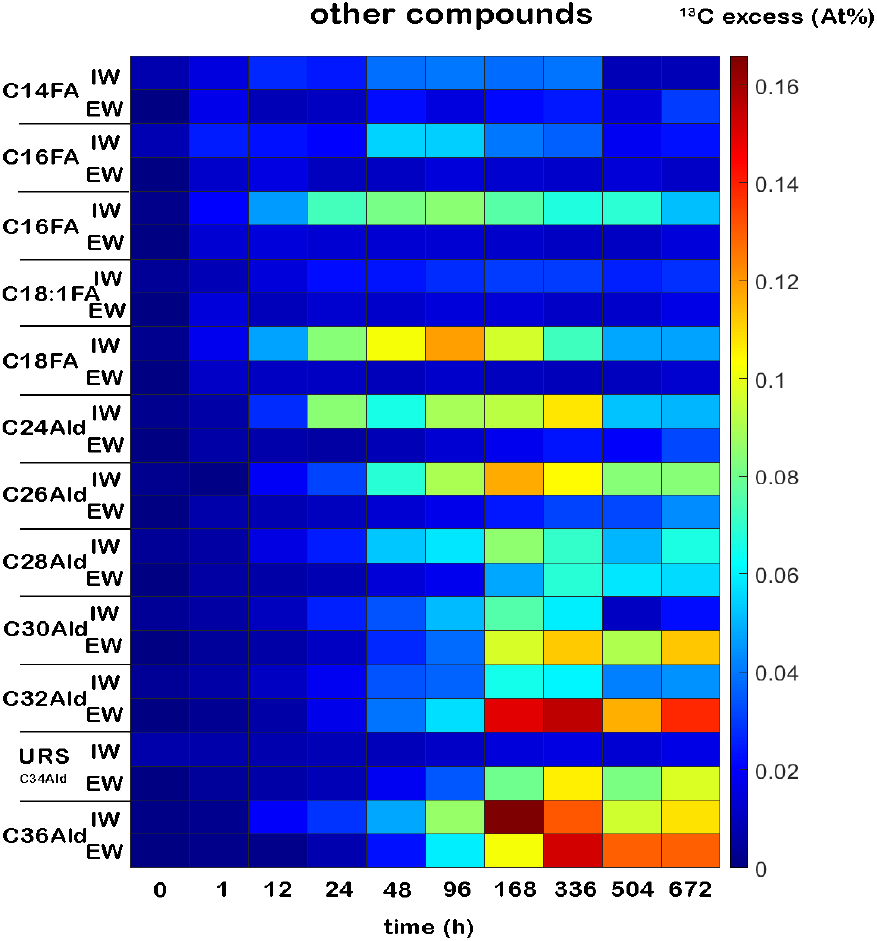
Time course of ^13^C excess of other compound than alkanes in the *P. laurocerasus* leaf cuticles separately for intra- (IW) and epicuticular wax (EW). Abbrevs. mean C14FA: Myristic acid, C16FA: Palmitic (native and TMS), C18:1FA: Oleic Acid, C18FA: Stearic acid, Ald: aldehyde, URS: ursolic acid, Cxx: number of carbon atoms in chain. Both leaf sides and treatments are pooled. N=8-32.

Other compounds detected were fatty acids and aldehydes (**SFig. 3, Fig. 7**). Short fatty acids (C16 and C18) were detected predominantly in the IW, however, their origin was presumably from the tissue (adsorbing to the cuticle during isolation as found previously). Aldehydes (present in both IW and EW), in turn, exhibited an interesting pattern. The shorter ones (C24, C26 and C28) were substantially enriched in IW, but the enrichment did not appear or was only marginal in EW. The longer ones (C30 and C32) showed the opposite behaviour with, significant enrichment only in the EW (**Fig. 7**).

To obtain comparison with annual plant species, we performed similar (pilot) experiments with *Brassica oleracea* (**SFig. 5**) and *Capsicum annuum* plants (**SFig. 6**). The common feature was a huge variability in ^13^C excess among plants and leaves, but leaf sides and wax compounds of particular leaf behaved very consistently (see discussion).

### Estimation of relative wax regeneration rates

Higher and faster EW ^13^C enrichment in collodion treated leaf-haves (EWF), compared to control ones (**Fig. 5, SFig. 4**) is (at least partly) due to reduced EW coverage after initial collodion treatment (mixing model with different pool sizes). This effect is stronger for adaxial leaf sides. For example, EW C31 alkane coverage was > 1 µg cm^-2^ in control ADAX leaf halves but only about 0.25 µg cm^-2^ in EWF ones at early times, partially converging towards the end of the experiment (**Fig. 3b**). Two approaches were applied to estimate the relative wax synthesis rate and equilibration of new wax between IW and EW.

We applied product of compound coverage multiplied by its ^13^C excess. It represents the relative amount of newly synthesized/deposited wax compound (Kubásek *et al*., 2023). This product indicates that EW deposition rate is slightly higher for the control than for the EWF leaf halves, but only on the ADAX sides (**Fig. 8a,b**). The behaviour of both, C29 (**Fig. 8a, c**) and C31 alkanes (**Fig. 8b, d**) is almost identical.

**Fig. 8.**
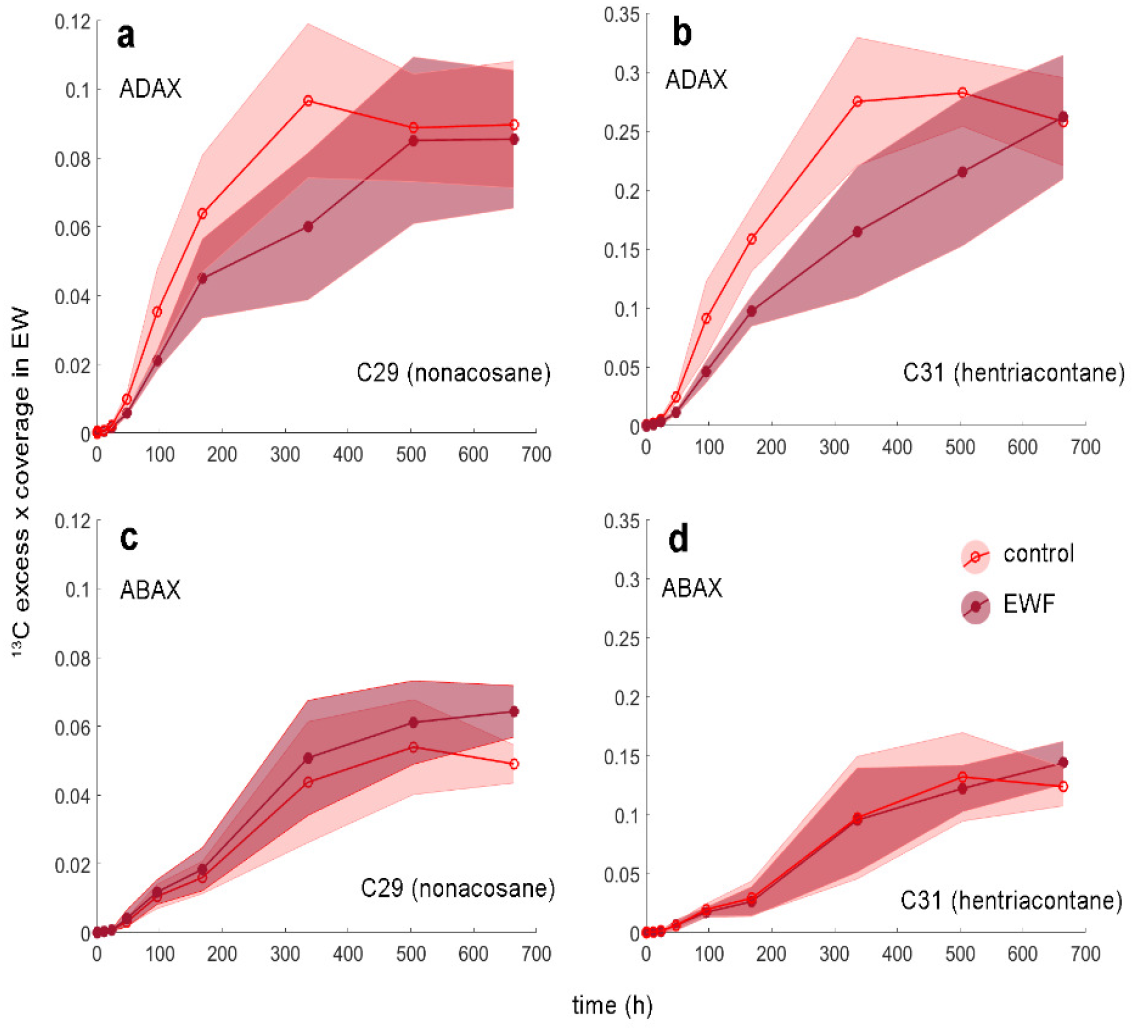
Estimation of the new epicuticular wax deposition as a product of particular compound ^13^C excess (At%) and its coverage (µg cm^-2^). This deposition is shown for two dominant alkanes (C29 n- nonacosane, **a,c**; and n-C31 hentriacontane, **b,d**), separately for adaxial (**a,b**) and abaxial (**c,d**) leaf side. Red lines and open symbols represent means for control leaf halves (control), purple lines and closed symbols represent means for leaf halves from which epicuticular wax was once partially removed at the start of the experiment (‘epicuticular wax free’, EWF). Shaded areas are one standard deviation. N=4.

Having all, IW and EW coverages and IW and EW ^13^C excess, we can calculate this product separately for IW and EW and estimate the fraction of new carbon in each layer (expressed as a “percentage of new wax in EW”). The values lead to the conclusion that about 60-80 % of the total new wax (^13^C enriched) is in EW towards the end of the experiment. Neither leaf side, nor control vs. EWF treatment, nor C29 vs. C31 alkane have substantial effect on the equilibration kinetics (**Fig. 9**; times up to ≈48 h are often too noisy for correct calculation due to negligible ^13^C excess in EW during these early times).

**Fig. 9.**
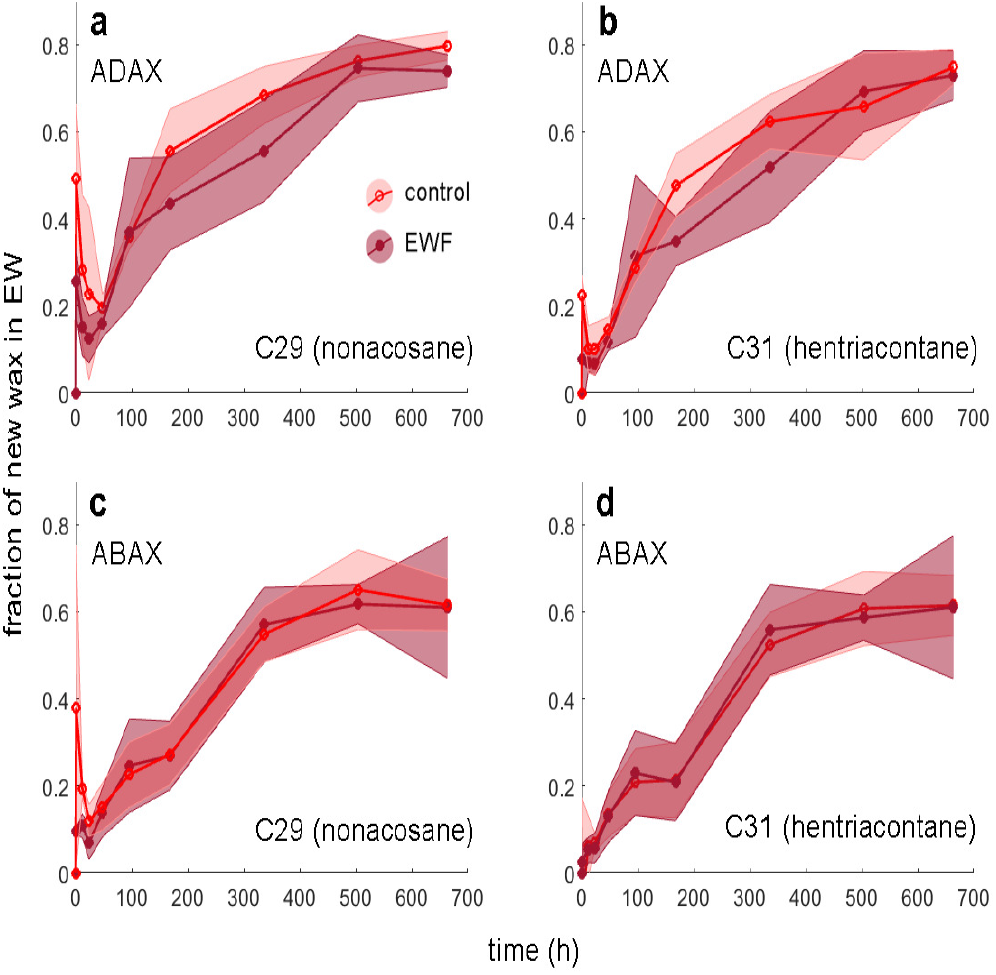
Fraction of the new wax deposition in epicuticular wax (EW) at particular times. This fraction is calculated as new wax coverage in EW divided by total coverage of new wax (IW+EW). The fraction is shown for two dominant alkanes (C29 nonacosane, **a,c**; and C31 hentriacontane, **b,d**), separately for adaxial (**a,b**) and abaxial (**c,d**) leaf sides. Red lines and open symbols represent means for control leaf halves (control), purple lines and closed symbols represent means for leaf halves from which epicuticular wax was once partially removed at the start of the experiment (‘epicuticular wax free’, EWF). Shaded areas are one standard deviation. N=4.

On the other hand, the turnover of ursolic acid (and oleanolic acid) is certainly very slow and difficult to estimate. This is the consequence of both, low ^13^C enrichment (**Fig. 7, STab. 1**) and high coverage, particularly in ursolic acid (**SFig. 3, STab. 1**). For other compounds, similar estimation is difficult due to their low coverage and/or possible GC co-elution with other compounds (retention time is the only identification method for GC-IRMS).

## Discussion

### Dominant alkanes, unlike triterpenoids, are renewed in both IW and EW

Consistently with the previous study on *Clusia rosea*, we confirmed that mature leaves synthetize substantial amounts of n-alkanes and deposit them first in the IW, then equilibrate with the EW pool (Kubásek *et al*., 2023). Because of the same labelling design, we can compare the results obtained. The ^13^C excess in dominant alkanes of young *C. rosea* leaves was up to 1 At% in average, which is several times higher than our results on mature *P. laurocerasus* leaves. On the other hand, mature *C. rosea* leaves reached about 0.2 At% ^13^C excess, which is pretty similar to the results in this study (**Fig. 5, SFig. 4**). This variable depends on the amount of wax previously deposited (dilution effect). It is therefore not a measure of the net wax synthesis rate but represents a nice proxy of wax turnover (wax turnover of young leaves seems to be several fold faster than mature leaves due to their lower wax coverage).

Pentacyclic triterpenoids reveal a different pattern. Ursolic acid is dominant in IW of *P. laurocerasus* (**SFig. 3**). We present here that the ^13^C excess is close to zero in IW ursolic and oleanolic acids during four weeks after ^13^C labelling (**Fig. 6, Fig. 7, STab 1**), indicating very slow turnover of these compounds. Some enrichment in EW ursolic acid is most likely chromatographic artefact (co-elution with C34 aldehyde, see Results).

We can speculate that the very high boiling point of triterpenoids (volatility is reciprocal to retention index), together with their huge coverage and IW localisation, protects them from the erosion/evaporation. Therefore, the new triterpenoids do not need to be regenerated or only minimally in mature leaves.

### Initial EW removal has marginal or no effect on new EW wax deposition rate

Cuticle formation impose substantial metabolic cost on plants (Onoda *et al*., 2012). Thus, the timing of cuticle ontogeny and its renewal under various stresses is certainly under evolutionary pressure, but remains poorly understood. Moreover, the multivariate optimum seems to be different for plants/leaves of various functional groups and leaf longevity (Neinhuis *et al*., 2001). An elegant atomic force microscopy (AFM) study, revealed quick regeneration of epicuticular wax films/crystals in leaves of various plant species after wax disturption (Koch *et al*., 2004). However, it was not known whether de-novo wax synthesis or so called ‘steam distillation’ of pre-existing intracuticular wax is responsible for epicuticular wax delivery to the mature leaf surfaces (Neinhuis *et al*., 2001). Kubásek et al. (2023) demonstrated that new wax synthesis and deposition occur at comparable rates in both, young and mature evergreen leaves of *Clusia rosea*. Here we extend this approach to the effect of EW wax removal, not addressed until now, for *P. laurocerasus*; another species with long leaf longevity. Initial collodion EW removal (‘epicuticular wax free’, ‘EWF’ treatment) had no effect on further EW deposition on lower leaf side (**Fig. 8c,d**), but seems to be slightly inhibitory on the upper side (**Fig. 8a,b**).

However, this method relies on the assumption that EW loss due to erosion/abrasion is negligible or comparable in all treatments during the experiment. This will be less important for the lower leaf side (ABAX), where the collodion EWF treatment removes lower fraction of EW (**Fig. 3c,d**), while further EW yield on the upper leaf side (ADAX) is down to 20 % of the initial coverage at the beginning and does not recover completely for the entire experiment (**Fig. 3 a,b**). Thus, any particular EW loss after EWF treatment will lead to an underestimation of the wax deposition rate, and the ADAX leaf side of *P. laurocerasus* will be more prone to this underestimation.

It is also worth noting that the EWF treatment was carried out using collodion, a nitrocellulose solution in organic solvents, which is suspected of not being selective enough for EW and/or of penetrating into the leaf tissue affecting its physiology (Jetter *et al*., 2000). We found that collodion is highly EW selective at least for leaves of *C. rosea* (Kubásek *et al*., 2023) and *P. laurocerasus* (this study). And, in agreement with recent work (Jetter & Riederer, 2016), it does not inhibit the leaf ability to produce new wax. We also checked whether the rate of photosynthesis (*An*) had changed by side-specific gasometric measurements (2x Li-Cor 6400 XT). *An* was not affected by EWF treatment on the stomatous ABAX leaf side and *An* was not measurable on the astomatous ADAX side of the leaves (data not shown).

### Newly deposited IW alkanes rapidly equilibrate with EW alkanes and this rate is unaffected by the EWF treatment

Our method is able to provide a completely new result - the equilibration rate of newly deposited IW compounds with EW compounds already present on the cuticle. **Fig. 9** shows that it takes approximately 250 to 350 h for the same amount of newly synthesised alkanes to be in IW and EW, respectively. Typically, 60 to 80 % of these are in EW at the experiment end (four weeks after labelling, ≈ 700 h). Surprisingly, neither EWF treatment nor leaf side has any effect on this kinetic for two dominant alkanes. It’s now clear that relatively immobile and high-boiling waxes (alkanes in this case) rapidly equilibrate between cuticle substructures, and the driving force for this may be water cuticular transpiration, called ‘steam distillation’ *sensu* Neinhuis (Neinhuis *et al*., 2001). It remains to be elucidated why triterpenoids do not follow this rule, being arrested largely in IW of many plant species (Buschhaus & Jetter, 2011).

### Minor compounds follow unprecedented pattern of ^13^C enrichment

Preferential ^13^C enrichment of IW short aldehydes (C24, C26, C28) and EW longer aldehydes (C30 and C32) is difficult to explain (**Fig. 7**). First, the C30 aldehyde should be a precursor of the dominant C29 alkane and the C32 aldehyde of the dominant C31 alkane (Bernard & Joubès, 2013). Therefore, one would expect the enrichment pattern for these pairs to be similar. No such similarity is evident when comparing **Fig. 6** and **Fig. 7**. Second, IW and EW are subjected to different environmental stresses that can lead to wax loss, compound interconversion and chemical degradation. On the other hand, most studies found that only very high, non-physiological, doses of UV-radiation, ozone and/or other oxidising agents led to substantial chemical changes in cuticle wax mixtures, but the thin outermost layer of the cuticle can be modified even under physiological conditions (for a comprehensive review see (Jetter *et al*., 2018). Third, the composition of P. *laurocerasus* IW and EW changed only ontogenetically in intact leaves, not in isolated cuticle or waxes(Jetter & Schäffer, 2001a). This suggest that rather than spontaneous chemical reactions, new wax synthesis and probably transport of wax compounds back into the tissue is likely. This topic certainly deserves further experimental work with high spatial-temporal resolution.

### There are contrasting differences between annual and long-lived leaves

*C. rosea* and *P. laurocerasus*, both species with evergreen leaves, studied so far using this method, showed surprisingly little variability among replicates, considering that four individual plants were subsequently labelled (where conditions may vary somewhat) and that leaves of slightly different ages had to be sampled. This leads us to conclude that wax renewal is uniform for similar species.

In contrast, the most striking result in our pilot experiments was the enormous variability even among very similar annual leaves. (**SFig. 5, SFig. 6**). For example, *Brassica oleracea* plants grown in a greenhouse sampled at 139 h revealed almost no enrichment despite substantial enrichments in neighbouring times. We can rule out a technical error, as both leaf sides and all wax compounds behave in the same way. We can speculate that this is a widespread economic and ecological strategy to stop wax synthesis in mature annual leaves at a certain time before senescence starts. The cuticle, even without maintenance, may function well for many days or even weeks after this ‘strategic stop’. This can save the plant’s carbon/energy resources for any other uses.

In conclusion, we confirmed continuous cuticular wax deposition in mature evergreen leaves of cherry laurel. We tested for the first time the effect of epicuticular wax (EW) removal on further wax synthesis/deposition, which appears to be marginal if any. We also found that intracuticular alkanes rapidly equilibrate with epicuticular alkanes and that the turnover of pentacyclic triterpenoids is surprisingly slow, results not reported before. Finally, consistent differences were found between long living and annual leaves.

## Acknowledgement

We are indebted to Jiří Šantrůček for the initial idea for this experiment and valuable comments on the manuscript, and to Petra Fialová for technical assistance.

