## Supplementary information for "Removal of epicuticular wax has no effect on the rate of cuticular wax deposition, which is constant in mature cherry laurel leaves"

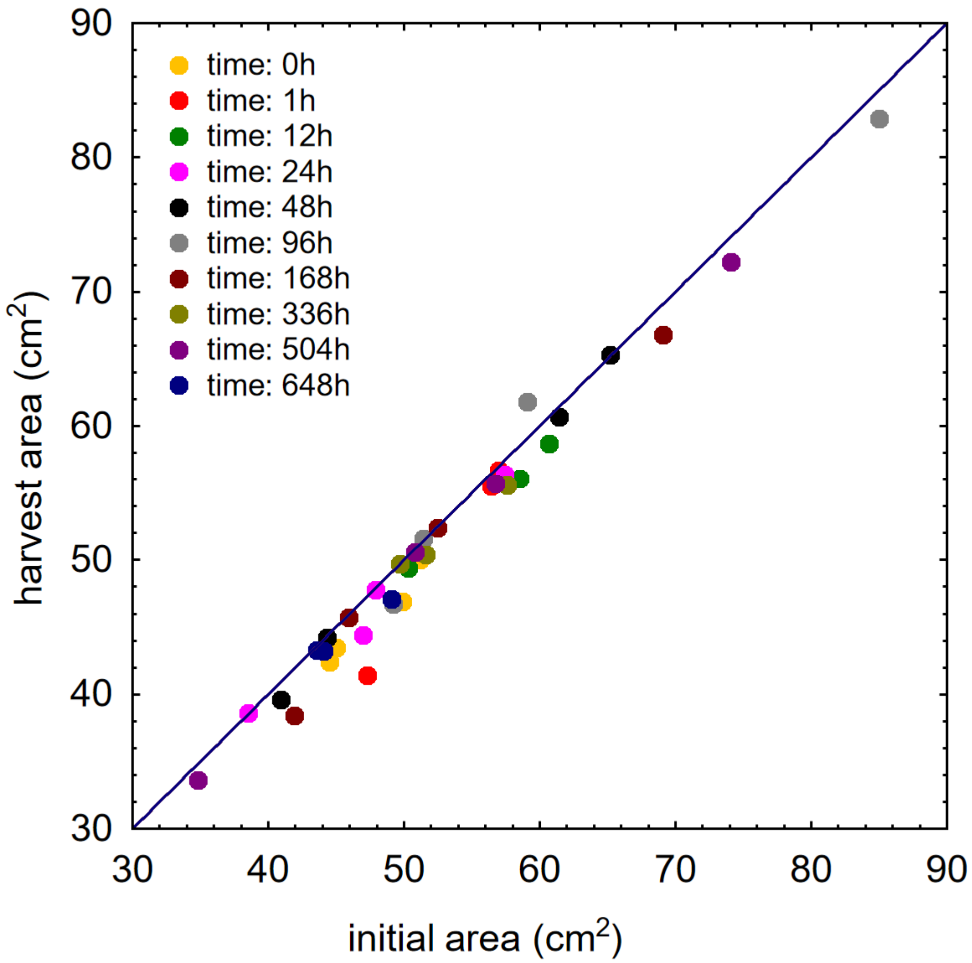


**SFig. 1. Correlation of initial and harvested leaf area of *P. laurocerasus*.** Initial areas were obtained before the experiment by drawing leaf margins on the paper, final areas were obtained at particular times (0 to 648h) by scanning harvested leaves with a scale bar. Areas were calculated using ImageJ software. The blue line indicates a 1:1 ratio.

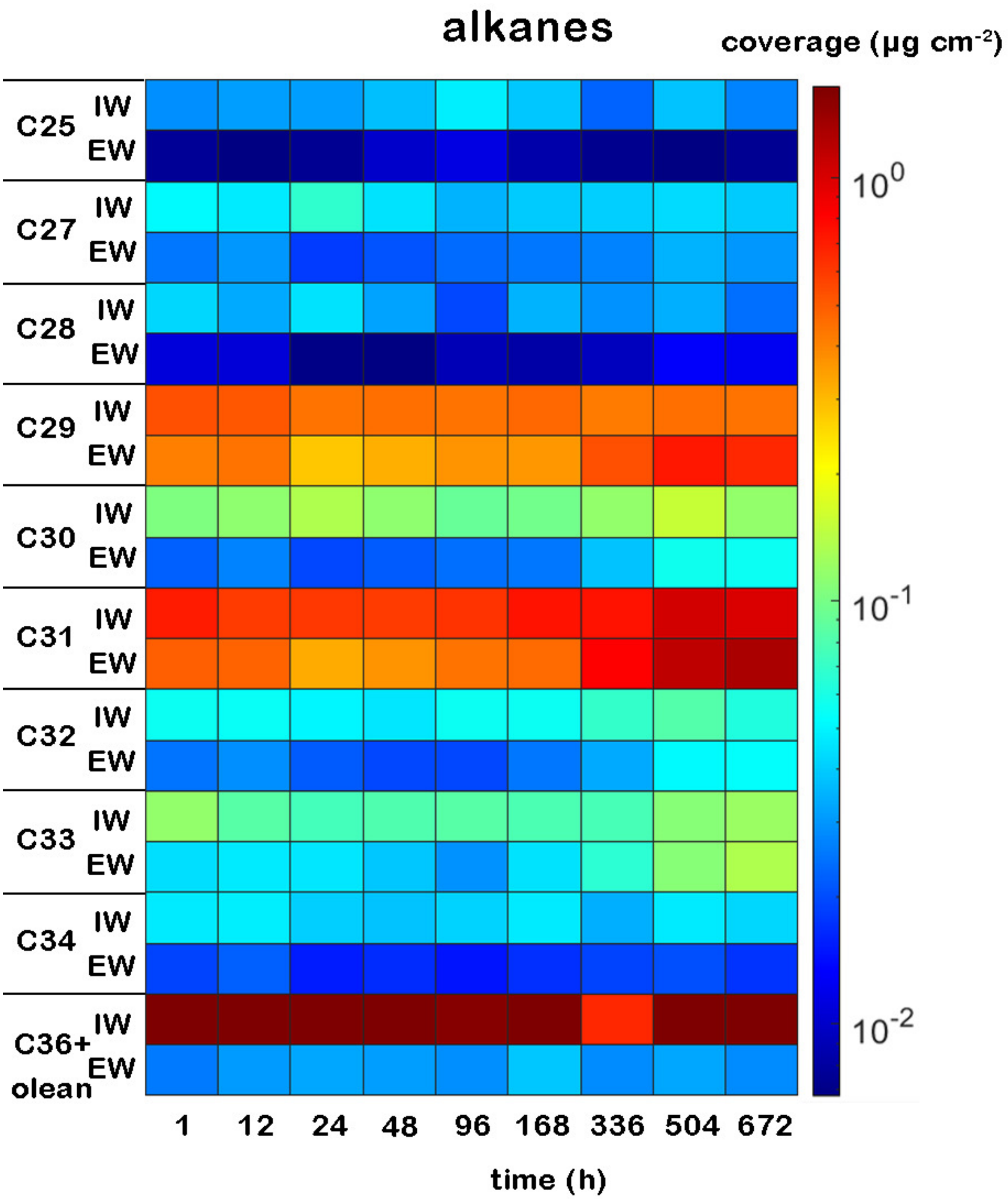


**SFig. 2. Coverage of n-alkanes detected in *P. laurocerasus* leaf cuticles separately for intra- (IW) and epicuticular wax (EW).** Oleanolic acid has retention index very close to that of alkane 36, therefore it co-eluted in the same peak and dominates in IW. Adaxial and abaxial sides and both treatments are pooled. Note that the colour scale is logarithmic. N=8-32.

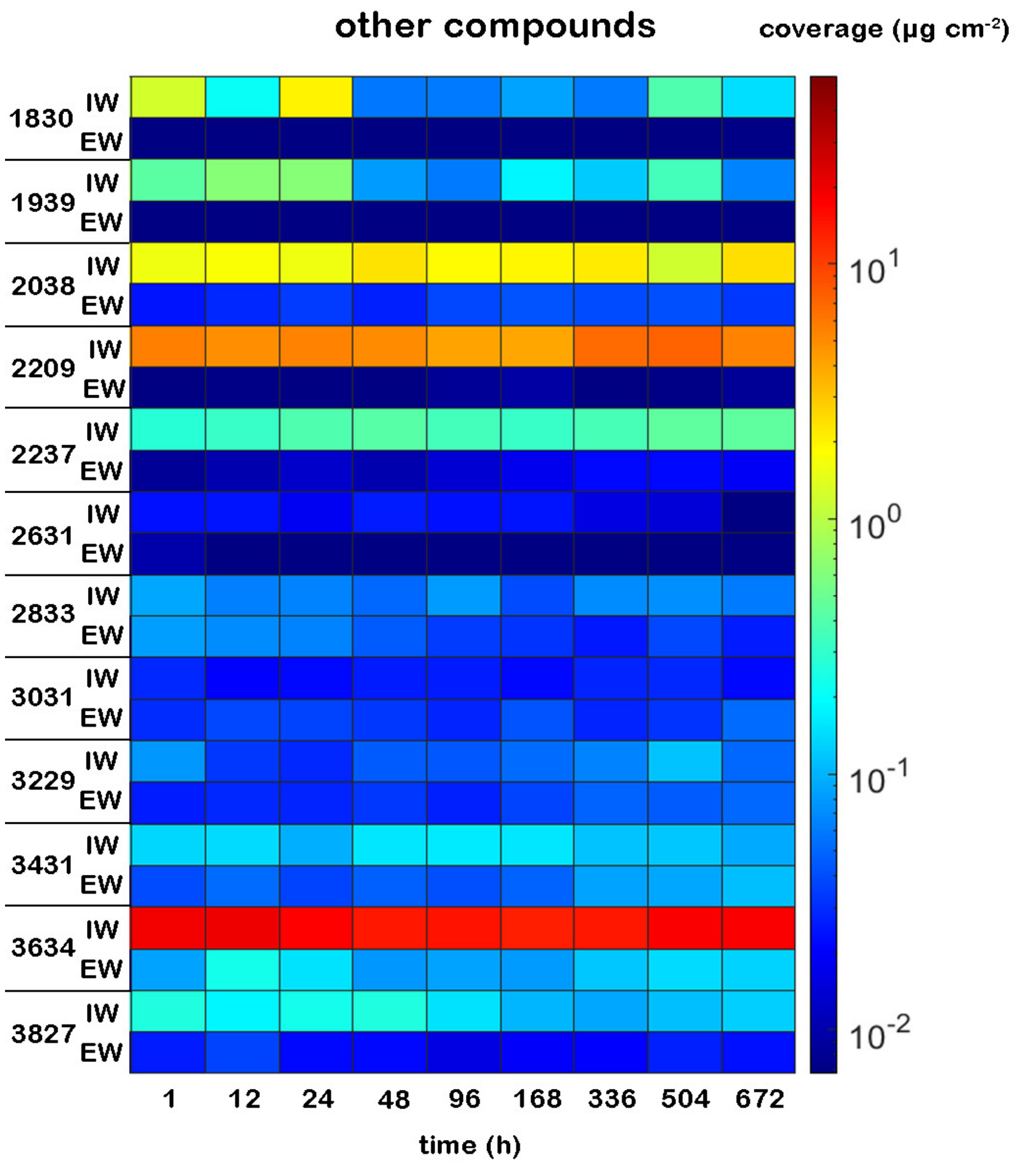


**SFig. 3. Coverage of other compounds than alkanes detected in *P. laurocerasus* leaf cuticles separately for intra- (IW) and epicuticular wax (EW).** Retention indices mean - 1830: Myristic acid, 1939: Palmitic acid-native, 2038: Palmitic acid, 2209: Oleic Acid, 2237: Stearic acid, 2631: C24 aldehyde, 2833: C26 aldehyde, 3031: C28 aldehyde, 3229: C30 aldehyde, 3431: C32 aldehyde, 3634: Ursolic acid and C34 aldehyde, 3827: C36 aldehyde. Adaxial and abaxial sides and both treatments are pooled. Note that the colour scale is logarithmic. N=8-32.

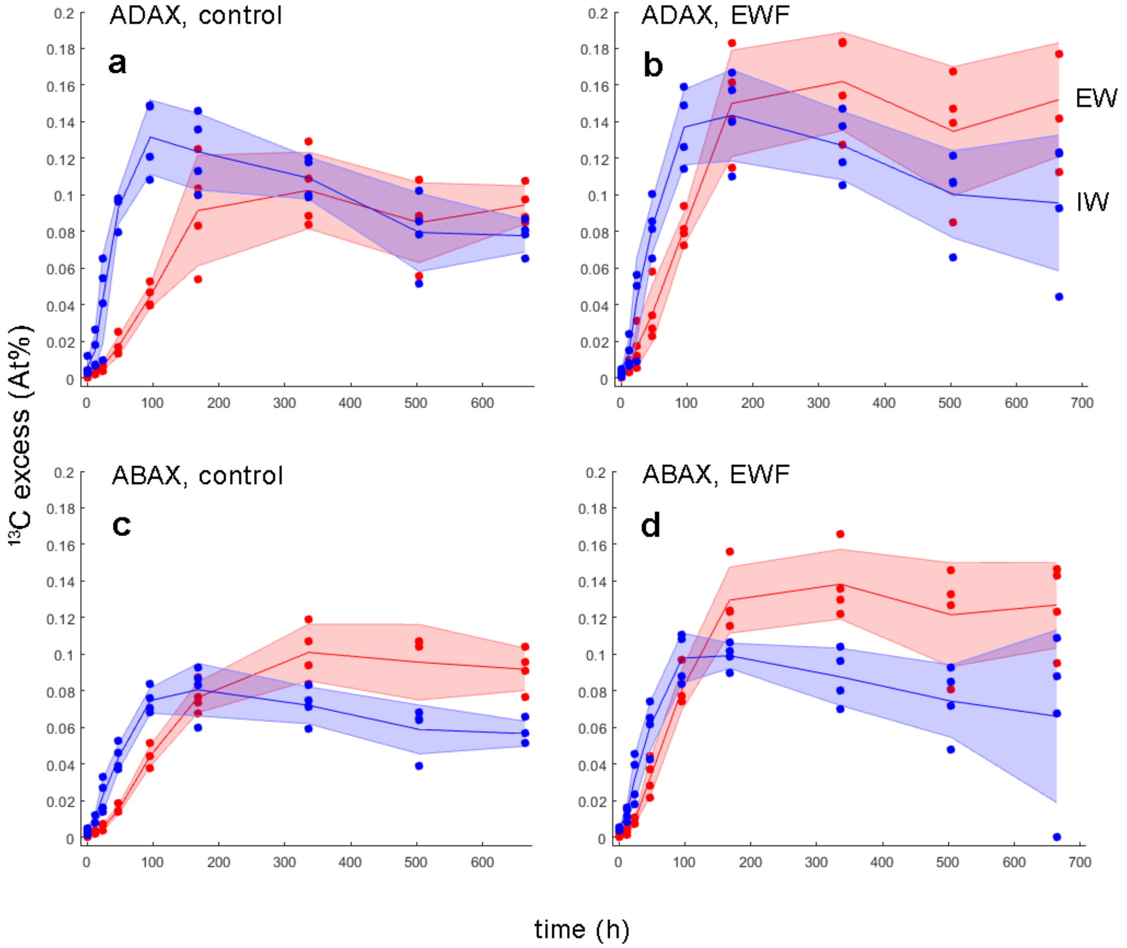


**SFig. 4. Timecourse of $^{13}C$ excess of n-nonacosane in the *P. laurocerasus* leaf cuticles.** Intracuticular wax (IW, blue) and epicuticular wax (EW, red) are shown separately. **a**,**b** show adaxial leaf side, **c**,**d** abaxial leaf side. **a**,**c** show unaffected leaf halves (control), **b**,**d** halves in which epicuticular wax was partially removed by collodion at the start of the experiment ('epicuticular wax free', EWF). Lines represent mean, shaded areas are one standard deviation N=4.

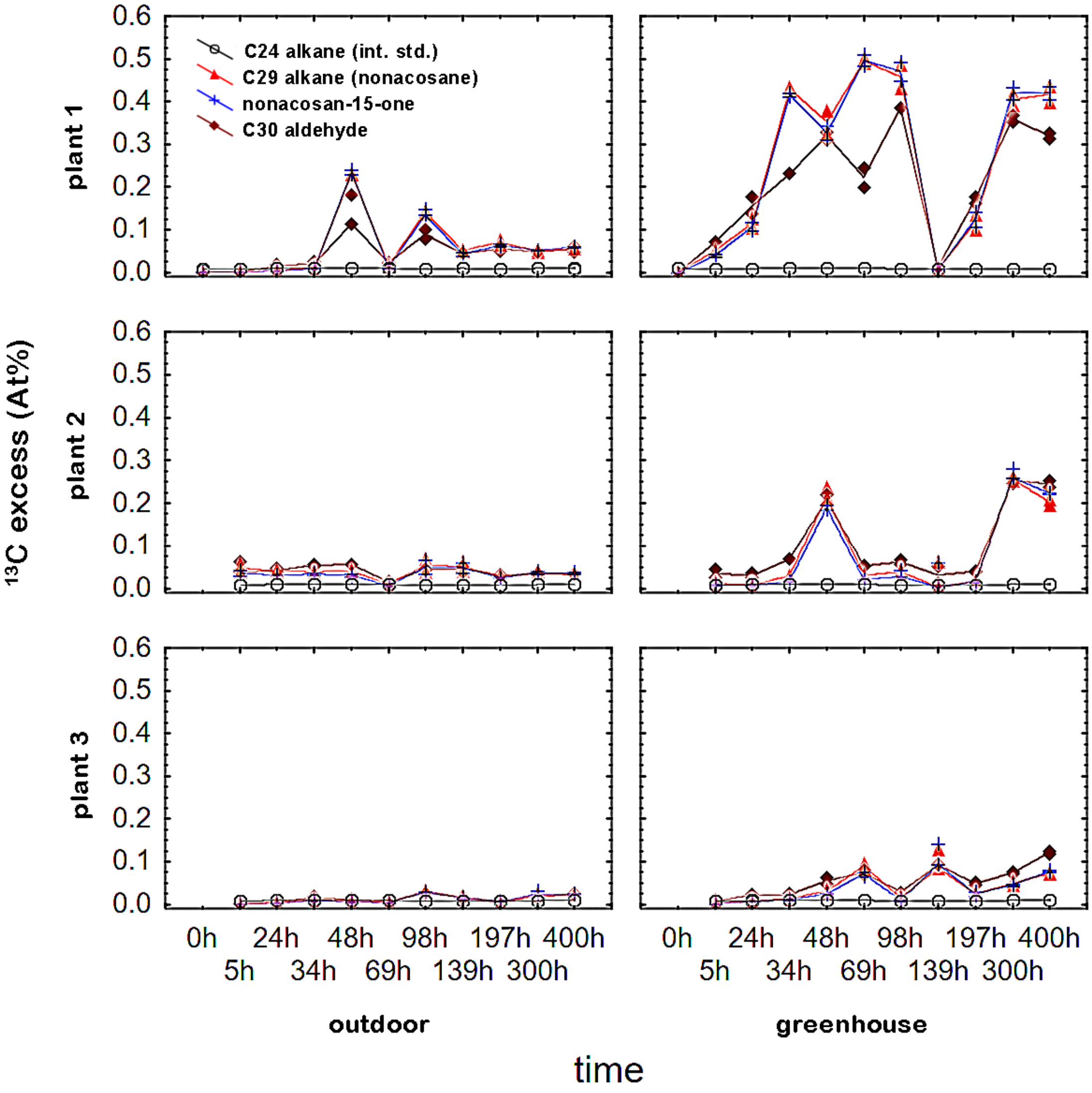


**SFig 5. Timecourse of $^{13}$C excess of three dominant compounds in epicuticular wax (EW) of *Brassica oleracea* var. italica leaves during our pilot experiment.** Plants were labelled with $^{13}$C and EW was sampled in the same way as in the main experiment with *P. laurocerasus*. No treatment was applied prior to labelling. After labelling, the plants were divided into two groups. One group was placed in a greenhouse with a controlled environment, similar to the *P. laurocerasus* experiment (see Material and methods) and labelled as 'greenhouse'. The second group, labelled 'outdoor', was placed on an outdoor terrace with a much more challenging environment (hot summer, with temperatures ranging from 15 °C at nights to over 33 °C on the hottest days). EW from the adaxial and abaxial leaf side were harvested separately and two identical symbols are shown (they are very close or overlap in most cases). There is a high variability rather than a consistent pattern. Therefore individual plants/values are shown, not descriptive statistics.

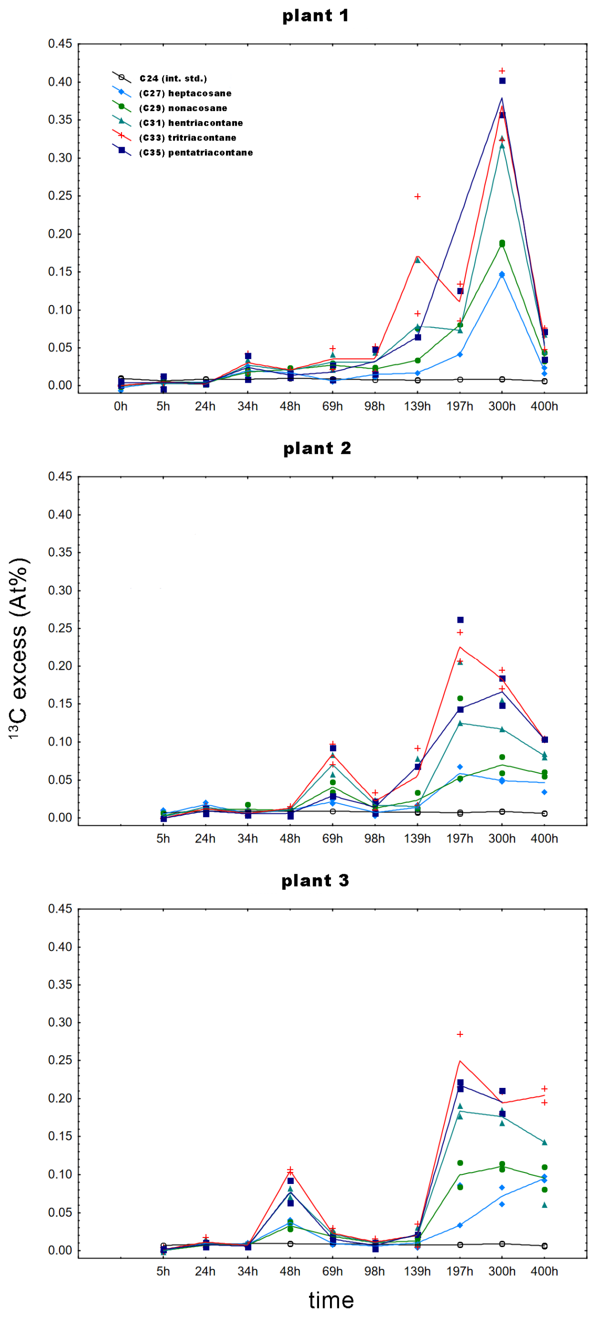


**SFig 6. Timecourse of $^{13}C$ excess of five alkanes in epicuticular wax of *Capsicum annum* leaves during our pilot experiment.** Plants were labelled with $^{13}C$ and EW sampled in the same way as in the main experiment on *P. laurocerasus*. No treatment was applied before the labelling. Plants were placed in the glasshouse with controlled environment similar to the *P. laurocerasus* experiment (see Material and methods). EW from the adaxial and abaxial leaf side were harvested separately and two identical symbols are shown (they are very close or overlap in most cases). There is a high variability rather than a consistent pattern. Therefore individual plants/values are shown, not descriptive statistics.

**STab. 1.** Ursolic acid coverage and $^{13}C$ excess (mean ± StD), separately for intra- (IW) and epicuticular wax (EW), and separately for leaf side and collodion treatments separately. IW/EW coverage stands for IW/EW coverage ratio, N=4.

| time | Treat | IW (µg cm-2) | EW (µg cm-2) | IW/EW coverage | IW 13C excess (At%) | EW 13C excess (At%) |
|---|---|---|---|---|---|---|
| 0h | CONT_U | 27.14 ± 2.2 | | | 0.002 ± 0 | |
| 0h | EWF_U | | | | | |
| 0h | CONT_L | 12.93 ± 3.04 | | | -0.001 ± 0.001 | |
| 0h | EWF_L | | | | | |
| 1h | CONT_U | 23.29 ± 13.02 | 0.12 ± 0.05 | 199 | 0.001 ± 0.001 | 0.002 ± 0.001 |
| 1h | EWF_U | 25.86 ± 5.35 | 0.04 ± 0.02 | 581 | 0.001 ± 0.001 | 0.005 ± 0.001 |
| 1h | CONT_L | 12.63 ± 6.99 | 0.14 ± 0.15 | 92 | -0.002 ± 0.001 | 0.005 ± 0.001 |
| 1h | EWF_L | 14.55 ± 3.02 | 0.05 ± 0.03 | 304 | -0.001 ± 0.001 | 0.005 ± 0.001 |
| 12h | CONT_U | 27.66 ± 3.94 | 0.14 ± 0.07 | 192 | 0.002 ± 0 | 0.004 ± 0 |
| 12h | EWF_U | 26.79 ± 2.14 | 0.09 ± 0.06 | 302 | 0.001 ± 0 | 0.008 ± 0 |
| 12h | CONT_L | 12.59 ± 4.04 | 0.45 ± 0.34 | 28 | 0 ± 0 | 0.006 ± 0 |
| 12h | EWF_L | 13.99 ± 3.64 | 0.22 ± 0.07 | 63 | -0.001 ± 0.001 | 0.006 ± 0.001 |
| 24h | CONT_U | 21.97 ± 3.49 | 0.09 ± 0.05 | 244 | 0.002 ± 0.002 | 0.01 ± 0.002 |
| 24h | EWF_U | 22.65 ± 4.13 | 0.1 ± 0.08 | 237 | 0.002 ± 0.001 | 0.01 ± 0.001 |
| 24h | CONT_L | 12.56 ± 4.06 | 0.28 ± 0.21 | 46 | 0.001 ± 0.002 | 0.007 ± 0.002 |
| 24h | EWF_L | 13.42 ± 4.05 | 0.15 ± 0.14 | 89 | 0.001 ± 0.002 | 0.006 ± 0.002 |
| 48h | CONT_U | 22.82 ± 15.62 | 0.1 ± 0.03 | 220 | 0.002 ± 0.002 | 0.023 ± 0.002 |
| 48h | EWF_U | 17.63 ± 10.54 | 0.05 ± 0.02 | 365 | 0.001 ± 0.001 | 0.031 ± 0.001 |
| 48h | CONT_L | 7.34 ± 10.07 | 0.12 ± 0.04 | 63 | 0.004 ± 0.001 | 0.01 ± 0.001 |
| 48h | EWF_L | 7.9 ± 4.45 | 0.04 ± 0.01 | 206 | 0.003 ± 0.001 | 0.01 ± 0.001 |
| 96h | CONT_U | 22.63 ± 1.98 | 0.09 ± 0 | 253 | 0.003 ± 0.002 | 0.052 ± 0.002 |
| 96h | EWF_U | 17.33 ± 14.37 | 0.05 ± 0.02 | 356 | 0.002 ± 0.001 | 0.059 ± 0.001 |
| 96h | CONT_L | 9.3 ± 3.29 | 0.12 ± 0.02 | 78 | 0.008 ± 0.001 | 0.013 ± 0.001 |
| 96h | EWF_L | 6.59 ± 4.53 | 0.09 ± 0.09 | 72 | 0.006 ± 0.001 | 0.014 ± 0.001 |
| 168h | CONT_U | 17.45 ± 12.65 | 0.1 ± 0.07 | 184 | 0.004 ± 0.002 | 0.127 ± 0.002 |
| 168h | EWF_U | 18.76 ± 15.28 | 0.04 ± 0.01 | 484 | 0.004 ± 0.003 | 0.161 ± 0.003 |
| 168h | CONT_L | 8.73 ± 5.79 | 0.1 ± 0.03 | 89 | 0.012 ± 0.005 | 0.022 ± 0.005 |
| 168h | EWF_L | 8.4 ± 6.62 | 0.07 ± 0.06 | 114 | 0.011 ± 0.003 | 0.031 ± 0.003 |
| 336h | CONT_U | 21.53 ± 1.44 | 0.14 ± 0.03 | 158 | 0.004 ± 0.002 | 0.14 ± 0.002 |
| 336h | EWF_U | 15.66 ± 9.27 | 0.06 ± 0.02 | 262 | 0.002 ± 0.002 | 0.179 ± 0.002 |
| 336h | CONT_L | 9.46 ± 4.54 | 0.16 ± 0.14 | 59 | 0.017 ± 0.007 | 0.054 ± 0.007 |
| 336h | EWF_L | 9.64 ± 4.73 | 0.11 ± 0.06 | 85 | 0.013 ± 0.003 | 0.053 ± 0.003 |
| 504h | CONT_U | 22.64 ± 5.04 | 0.16 ± 0.01 | 143 | 0.002 ± 0.001 | 0.11 ± 0.001 |
| 504h | EWF_U | 19.34 ± 7.2 | 0.09 ± 0.05 | 224 | 0.002 ± 0.001 | 0.123 ± 0.001 |
| 504h | CONT_L | 15.49 ± 5.63 | 0.19 ± 0.14 | 82 | 0.011 ± 0.003 | 0.048 ± 0.003 |
| 504h | EWF_L | 14.73 ± 4.24 | 0.13 ± 0.05 | 110 | 0.01 ± 0.003 | 0.046 ± 0.003 |
| 648h | CONT_U | 19.77 ± 4.36 | 0.19 ± 0.08 | 105 | 0.004 ± 0.002 | 0.107 ± 0.002 |
| 648h | EWF_U | 24.52 ± 2.96 | 0.14 ± 0.01 | 175 | 0.004 ± 0.001 | 0.148 ± 0.001 |
| 648h | CONT_L | 16.1 ± 2.41 | 0.13 ± 0.04 | 124 | 0.015 ± 0.004 | 0.052 ± 0.004 |
| 648h | EWF_L | 14.58 ± 3.34 | 0.08 ± 0.03 | 189 | 0.013 ± 0.004 | 0.083 ± 0.004 |
